# Discovery and Characterisation of a Plant GH1 β-Glucosidase Exhibiting Hydrolytic Activity on an *N*-Linked Glucopyranoside

**DOI:** 10.64898/2025.12.17.694871

**Authors:** Hani Gharabli, Carlotta Chiesa, Maher Abou Hachem, Ditte Hededam Welner

## Abstract

β-Glucosidases (Bgls) catalyse the hydrolysis of the glycosidic bond of β-D-glycosides. Ubiquitous in nature, they play vital biological roles across diverse organisms, and their versatility has made them valuable for industrial applications. Bgls are commonly known to hydrolyse *O*- and *S*-linked glycopyranosides, but their activity on *N*-linked glycopyranosides has yet to be demonstrated. In a previous study, we discovered and biocatalytically produced a novel *N*-glucopyranoside, methyl anthranilate-*N*-β-D-glucopyranoside (MANT-*N*-glucose). Building on this, we sought to develop a biocatalytic method for the degradation of MANT-*N*-glucose into its two main components, methyl anthranilate and glucose. Through screening of a eukaryotic Bgl library, we identified ZmGlu1 as an enzyme capable of hydrolysing MANT-*N*-glucose. This reaction displayed substantially reduced catalytic efficiency (0.31 min^-1^ mM^-1^) relative to ZmGlu1’s activity with its native substrate and other *O*-glucopyranosides. Structural modelling of enzyme–substrate complexes revealed key interactions likely contributing to the reduced activity. These findings provide a foundation for future investigations into *N*-glycopyranoside Bgl reactivity and highlight the broader potential of Bgls in biocatalytic applications.

**Importance:** In this study, we uncover a previously unknown function within the widely used enzyme, β-glucosidases (Bgls). These enzymes catalyse the hydrolysis of *O*- and *S*-linked glycopyranosides, yet their ability to act on *N*-linked glycopyranosides has remained unrecognised until now. Using both human gut bacteria and a library of eukaryotic Bgls, we identified and characterised an enzyme capable of hydrolysing an *N*-linked substrate, methyl anthranilate-*N*-β-D-glucopyranoside. This finding expands the recognised activity range of Bgls and highlights their broader industrial relevance. In addition, this finding opens avenues for further investigation into Bgls substrate selectivity, as well as enzyme engineering aimed at enhancing activity toward *N*-linked glycopyranosides and elucidating the underlying structure–function.

## Introduction

β-Glucosidases (Bgls, E.C. 3.2.1.21) are enzymes that catalyse the hydrolysis of glycosidic bonds to release the non-reducing terminal glucosyl residue from glycosides or oligosaccharides (Cairns and Esen, 2010). Bgls belong to the enzyme class, glycoside hydrolases (GHs, E.C. 3.2.1.-), which are divided into families according to their sequence and structure similarity (Henrissat, 1991; Henrissat and Bairoch, 1993). Currently, 194 GH families are listed on the Carbohydrate-Active Enzyme (CAZy) database (12^th^ November 2025, http://www.cazy.org) (Drula et al., 2022). Bgls are distributed across several GH families, including GH1, GH2, GH3, GH5, GH16, GH30, GH39, GH116, GH131, GH175, and GH180. Among these, the GH1 family contains the largest proportion of Bgls.

Bgls are ubiquitous in nature and play crucial roles in processes such as biomass conversion, host-pathogen interactions, and phytohormone activation (Cairns and Esen, 2010; Jin et al., 2011; Srivastava et al., 2019). Their distinctive catalytic properties also make them valuable in industrial applications (Yang et al., 2024), including their use as additives in food production (Souto et al., 2023) and in the generation of biofuels (Mahapatra and Manian, 2022). GH1 Bgls commonly catalyse the hydrolysis of *O*-linked glycosides. However, a smaller subset of this enzyme family, known as myrosinases (E.C. 3.2.147), can also hydrolyse *S*-linked glycosides, such as glucosinolates (Pardini et al., 2021). To the best of our knowledge, no Bgl has been described to cleave small molecule *N*-linked glycopyranosides.

In a previous study, we biocatalytically produced a novel *N*-glucopyranoside, methyl anthranilate-*N*-β-D-glucopyranoside (MANT-*N*-glucose) (Gharabli et al., 2025). Building on this discovery, we aimed to identify a GH capable of hydrolysing this product. To achieve this, we pursued two approaches: (i) *in vivo* testing of MANT-*N*-glucose on intestinal bacteria known for glycoside metabolism and diverse carbohydrate-active enzymes (Kaoutari et al., 2013; Luis and Martens, 2018; Theilmann et al., 2017), and (ii) screening an in-house library of GH1 Bgls from eukaryotes. These efforts revealed a maize Bgl, ZmGlu1, which hydrolysed MANT-*N*-glucose to its corresponding aglycon and glucose. The reaction was characterised, while structural modelling of the ZmGlu1:MANT-*N*-glucose complex provided insights into its distinct kinetics compared with the hydrolysis of its natural substrate, DIMBOA-glucoside.

## Materials and methods

### Chemicals and reagents

MANT-*N*-glucose was produced in-house with a purity > 99 % (HPLC) (Gharabli et al., 2025). All other substrates and reagents were obtained from Sigma unless stated otherwise.

### HPLC analysis for MANT and MANT-*N*-glucose detection and quantification

Product formation was followed by HPLC, using an Ultimate 3000 Series apparatus (Thermo Fisher) employing a Kinetex C18 analytical column (dimensions 4.6 x 100 mm; pore size 100 Å; particle size 2.6 μm) (Phenomenex). 0.1 % formic acid in water (A) and acetonitrile (B) were used as mobile phases employing a gradient elution: 0.00–0.50 min, 2% B;0.50-0.51 min, linear ramp to 35 % B; 0.51–1.90 min, 35 % B; 1.90-2.50 min, linear ramp to 100 % B; 2.50–4.20min, 100 % B; 4.20-4.21, linear ramp to 2 % B; 4.21-5.00 min, 2 % B. The flow rate was set to 1 mL/min. The analytes were detected at 330 nm. The HPLC data were monitored and quantified via the Chromeleon software (Thermo Fisher Scientific). The product, MANT, was quantified using a calibration curve in all experiments except the analysis of MANT-*N*-glucose conversion in intestinal bacteria cultures and the kinetic characterisation. Here, the formation of the product was determined by analysing the ratio between the product peak and acceptor peak on the HPLC chromatograms, assuming the absorbance at 330 nm is equal.

### Screening intestinal bacteria for MANT-*N*-glucose degradation

Seven bacterial species isolated from the human gut were selected to screen for MANT-*N*-glucose degradation, assuming it would support strain growth. This included strains, *Bacteroides ovatus* DSM1896, *B. thetaiotaomicron* DSM2079, *Bifidobacterium longum* subsp. *longum* 1-6B, *Blautia hansenii* DSM20583, *Eubacterium ramulus* DSM15684, *E. ramulus* DSM16296, and *Roseburia intestinalis* DSM14610. These strains were grown at 37 °C without agitation in an anaerobic cabinet (Coy Laboratory Products, USA) under an 85 % N_2_/10 % H_2_/5 % CO_2_ atmosphere. *Blautia, Roseburia*, and *Bifidobacterium* strains were grown in YCFA (yeast extract-casein hydrolysate-fatty acids) medium (Duncan et al., 2002), while *Bacteroides* and *Eubacterium* strains were grown in CFA (YCFA lacking the yeast extract) medium to minimise growth on yeast extract. All were supplemented with 0.5 % (w/v) glucose. Growth experiments were performed by diluting 1 % (v/v) of an overnight culture (2 mL) into YCFA/CFA media containing either 0.5 % (w/v) MANT-*N*-glucose or 0.5 % (w/v) glucose. A negative control containing no carbon source was also included for each bacterial strain. 200 µL cultures were incubated in 96-well plates sealed with a Breathe-Easy® (Sigma-Aldrich, USA) sealing membrane at 37 °C without agitation in an Epoch2 microplate spectrophotometer (BioTek, Agilent, USA). Bacterial growth was monitored every 30 minutes for 24 hours by measuring the OD_600_. The data were analysed as means ± standard deviations of four replicate experiments.

### Analysis of MANT-*N*-glucose degradation in the supernatant of *B. ovatus* DSM1896 and *E. ramulus* DSM15684 cultures

The supernatants of *B. ovatus* DSM1896 and *E. ramulus* DSM15684 cultures, grown with MANT-*N*-glucose as the carbon source, were analysed over 36 hours using HPLC. A 1 mL culture, inoculated with 1 % (v/v) overnight culture in CFA supplemented with 0.5 % (w/v) MANT-*N*-glucose, was incubated at 37 °C without agitation. A non-inoculated control was included in the experiment. All cultures were grown in duplicates. 50 µL samples were taken at 0, 8, 12, 16, 24, 32, and 36 hours. Samples were centrifuged, and the supernatant was isolated and stored at -80°C until further processing. For the analysis, the samples were thawed, and 25 µL of the sample was diluted in 25 µL water and 50 µL methanol. The diluted sample was then centrifuged, and 50 µL of the supernatant was transferred to 200 µL of water and subsequently transferred to the HPLC. The injection volume was set to 7 µL. A two-way ANOVA test was performed to determine statistical significance using a significance level of 0.05.

### Cloning and expression of Bgls

The full-length coding sequence of the Bgls was synthesised and cloned into the pET30a(+) vector by Genscript (USA) or Biomatik (USA), using the NcoI and XhoI restriction sites. The protein-coding sequences included an N-terminal 6xHis-tag and a TEV-cleavage site. Plasmids were transformed into *E. coli* BL21 Star™ (DE3) cells (Thermo Fisher Scientific). Precultures were prepared by inoculation from glycerol stocks in 2xYT medium supplemented with 50 μg mL^−1^ kanamycin and grown overnight at 37 °C, 250 rpm. Precultures were then diluted to OD_600_ = 0.05 in 2xYT supplemented with 50 μg mL^−1^ kanamycin in a baffled flask. The cultures were incubated at 37 °C, 180 rpm, until the OD_600_ reached 0.6-1.0, upon which it was induced with 0.2 mM IPTG and subsequently incubated overnight at 20 °C, 180 rpm. The cells were harvested by centrifugation at 4500 g, 30 min at 4 °C. The cell pellet was washed once with phosphate-buffered saline (PBS), pH 7.4. The isolated cell pellet was either stored at -70 °C or processed for lysis and purification.

### Preparation of Bgl cell lysates and screening of MANT-*N*-glucose hydrolysing activity

The Bgl cell lysates were prepared from the cell pellet of 50 mL cultures. The resulting pellet was resuspended in 10 mL storage buffer (50 mM phosphate, 150 mM NaCl, pH 7.5) with 10 μg/mL DNase. The cell suspension was lysed via sonication using a Sonics Vibra Cell (USA) at 70 % amplitude with 15 seconds on/30 seconds off intervals for a total of 2.5 minutes of sonication time. The soluble fraction was then isolated by centrifuging the lysed sample at 14,500 × g at 4 °C for 45 min, followed by filtration through a 0.45 µm syringe filter. The filtered lysates were then aliquoted and stored at -70°C.

The lysates were assayed against MANT-*N*-glucose to establish their respective hydrolytic activity. 40 µL of the lysate was mixed with 5 mM of MANT-*N*-glucose in 50 mM phosphate buffer, pH 8.0, to a final volume of 200 µL. The lysate of WT *E. coli* and a reaction mixture without lysate were included as controls, respectively. The reaction mixtures were incubated overnight at room temperature. The reaction mixtures were processed for HPLC analysis by diluting 1:1 in methanol, centrifuging the diluted samples and filtering the supernatant through a 0.22 µm syringe filter. The filtered sample was diluted 1:4 in water and transferred to the HPLC. The injection volume was set to 15 µL.

### Purification of *Zm*Glu1

The purified *Zm*Glu1 was prepared from the cell pellet of 1000 mL cultures. The resulting pellet was resuspended in an appropriate amount (10 mL pr. g of the pellet) of lysis buffer (50 mM phosphate, 300 mM NaCl, 20 mM imidazole, pH 7.5, supplemented with 10 μg/mL DNase, 0.1 mg/mL lysozyme, 0.05 % (v/v) Triton X-100). Cell lysis was performed by sonication on a Sonics Vibra Cell (USA) at 85 % amplitude with 15 seconds on/30 seconds off intervals for a total of 4 minutes of sonication time. The soluble cell lysate was recovered by centrifugation at 14,500 × g at 4 °C for 45 min. The supernatant containing the recombinant proteins was filtered with a 0.45 μm filter and purified by nickel-affinity chromatography (prepacked HisTrap™ FF columns, GE Healthcare) on an ÄKTA pure (GE Healthcare). Before sample loading, the column was washed and equilibrated with buffer A (50 mM phosphate, 300 mM NaCl, pH 7.5, with 20 mM imidazole). Following protein binding, the column was washed with buffer A and subsequently eluted using a gradient elution with buffer A and buffer B (50 mM phosphate, 300 mM NaCl, pH 7.5, with 500 mM imidazole). The elution peak fractions were pooled together and concentrated using centrifugal filters to a volume of 2.5 mL. The buffer was exchanged for the storage buffer (50 mM phosphate, 150 mM NaCl, pH 7.5) via PD10 desalting columns packed with Sephadex G-25 resin (Cytiva). The total concentration of the purified proteins was measured by spectrophotometric measurements at 280 nm using a NanoDrop 2000 (Thermo Fisher Scientific) and adjusted using the extinction coefficient. The sample purity was assessed using sodium dodecyl sulfate-polyacrylamide gel electrophoresis (SDS-PAGE). The purified protein was then aliquoted and stored at -70 °C.

### MANT calibration curve

Two calibration curves of MANT were prepared using a 5 mM stock in 20 % (v/v) DMSO in water. The calibration curves ranged from 100 to 10 µM and 500 to 50 µM, respectively, with 10 equally spaced concentrations diluted in water. The calibration curves met the following quality requirements: a correlation coefficient ≥ 0.99 and passage through (0, 0).

### Validation of lysate screening results with purified *Zm*Glu1

The hit achieved from screening the Bgl cell lysates against MANT-*N*-glucose was validated by assaying the purified *Zm*Glu1. The reaction mixture consisted of 3 µM of purified *Zm*Glu1 and 0.5 mM MANT-*N*-glucose dissolved in 50 mM citrate-phosphate buffer, pH 6.0, 50 mM phosphate buffer, pH 7.0, 50 mM phosphate buffer, pH 8.0, 50 mM glycine buffer, pH 9.0, or 50 mM glycine buffer, pH 10.0, respectively. A control was included for each condition without enzyme added to the mixture. The reaction mixtures were incubated overnight at 25 °C with agitation (500 rpm) on a heat block. The reaction mixtures were subsequently diluted 1:1 in methanol and centrifuged. 100 µL of the supernatant was transferred to 200 µL of water and analysed via HPLC. The injection volume was set to 15 µL.

### Identification of the optimal pH for *Zm*Glu1-catalysed MANT-*N*-glucose hydrolysis

Reactions were conducted over a pH range to determine the optimum pH for *Zm*Glu1-catalysed MANT-*N*-glucose hydrolysis. The following buffering agents were utilised for their respective pH ranges: pH 4.5-6.0, 50 mM citrate-phosphate; pH 6.0-8.0, 50 mM phosphate; pH 9-10, 50 mM glycine. Each reaction contained 1 mM MANT-*N*-glucose and 10 µM *Zm*Glu1. To account for the spontaneous degradation of MANT-*N*-glucose under acidic and basic conditions, controls were included in the experiment without the enzyme but with the addition of the equivalent volume of storage buffer for each tested parameter. The reaction mixtures were incubated at 25°C for 1 hour in a thermocycler. The reaction was terminated by thermal denaturation at 95 °C for 20 seconds. The reaction mixtures were then centrifuged, and 40 µL of the supernatant was diluted in 320 µL of 50 mM phosphate buffer, pH 8.0, to halt any spontaneous degradation before transferring to the HPLC. The injection volume was set to 25 µL. The data were analysed as means ± standard error of three biological replicates. The product peak areas from the controls were subtracted from those in the reaction mixtures containing the enzyme.

### Identification of the optimal reaction temperature for *Zm*Glu1-catalysed MANT-*N*-glucose hydrolysis

Reactions were performed at a temperature range to determine the optimum reaction temperature for *Zm*Glu1-catalysed MANT-*N*-glucose hydrolysis. Each reaction contained 1 mM MANT-*N*-glucose and 8 µM *Zm*Glu1 in 50 mM phosphate buffer, pH 6.0. To account for the spontaneous degradation of MANT-*N*-glucose, controls were included in the experiment without the enzyme, but with the addition of the equivalent volume of storage buffer for each tested parameter. The reaction mixtures were incubated at temperatures ranging from 30 °C to 54 °C in a thermocycler for 1 hour.

The reaction was terminated by thermal denaturation at 95 °C for 20 seconds. Subsequently, the reaction mixtures were centrifuged, and 40 µL of the supernatant was diluted in 320 µL of 50 mM phosphate buffer, pH 8.0, to prevent any spontaneous degradation before being transferred to the HPLC. The injection volume was set to 25 µL. The data were analysed as means ± standard error of three biological replicates. The product peak areas from the controls were subtracted from those in the reaction mixtures containing the enzyme.

### Kinetic characterisation of *Zm*Glu1-catalysed MANT-*N*-glucose hydrolysis

The kinetic properties of *Zm*Glu1-catalysed MANT-*N*-glucose hydrolysis were determined using the established optimal pH and temperature conditions. Reaction mixtures contained 3 µM *Zm*Glu1 and a MANT-*N*-glucose concentration ranging from 20 mM to 1.17 mM in 50 mM phosphate buffer, pH 6.0. The reaction mixtures were incubated at 35 °C on a thermocycler. The reaction was stopped by thermal denaturation at 95 °C for 20 seconds. The kinetic parameters were obtained by setting up two individual experiments where *Zm*Glu1 from two different purification batches were utilised, and two different sets of time points were employed. These were 5, 10, 15, 20 minutes and 10, 20, 30, 40 minutes, respectively. A control was included in each experiment without the addition of enzyme, but with the addition of an equivalent volume of storage buffer. A control was only included with the first time point since MANT-*N*-glucose is relatively stable in the employed conditions. The reaction mixtures were analysed via HPLC, and the data were analysed as means ± standard error. Michaelis-Menten plots were generated and analysed in Prism v10.4.0. The product peak areas from the controls were subtracted from those in the reaction mixtures containing the enzyme.

### Structural modelling and visualisation

A structural model of the *Zm*Glu1:MANT-*N*-glucose complex was generated using the prediction tool, Chai-1 (Boitreaud et al., 2024). The FASTA sequence of the protein and the SMILES of MANT-N-glucose were used as input, and the tool was run with MSAs enabled (JackHMMER). The predicted complex structure was visualised using PyMOL (v2.5.2). Intermolecular interactions were analysed using PLIP (protein ligand interaction profiler) (Adasme et al., 2021) and manual inspection of the complex structures.

## Results and discussion

### The metabolism of MANT-*N*-glucose by intestinal bacteria

A panel of seven intestinal bacteria was screened against MANT-*N*-glucose, with the assumption that its degradation would release glucose and promote growth. Most species showed some growth, with *B. ovatus* DSM2096 and *E. ramulus* DSM15684 exhibiting the greatest increase in cell density (Figure 1). These strains were selected for further testing to assess the correlation between growth and MANT-*N*-glucose conversion. Over 36 hours, *E. ramulus* cultures showed no significant conversion compared to the control (P = 0.6294), whereas *B. ovatus* cultures displayed a significant increase (P < 0.0001), indicating a link between cell density and MANT-*N*-glucose conversion (Supplementary Figure 1). However, a depletion of the glycoside was not observed (Supplementary Figure 1), as previously reported in a similar experiment (Theilmann et al., 2017). Given this outcome and positive *in vitro* Bgl screening results (*vide infra*), the phenomenon was not pursued further, though an *N*-glucopyranoside-hydrolysing mechanism in *Bacteroides* remains plausible, potentially involving an orchestra of glucosidases (Nasseri et al., 2024).

**Figure 1.**
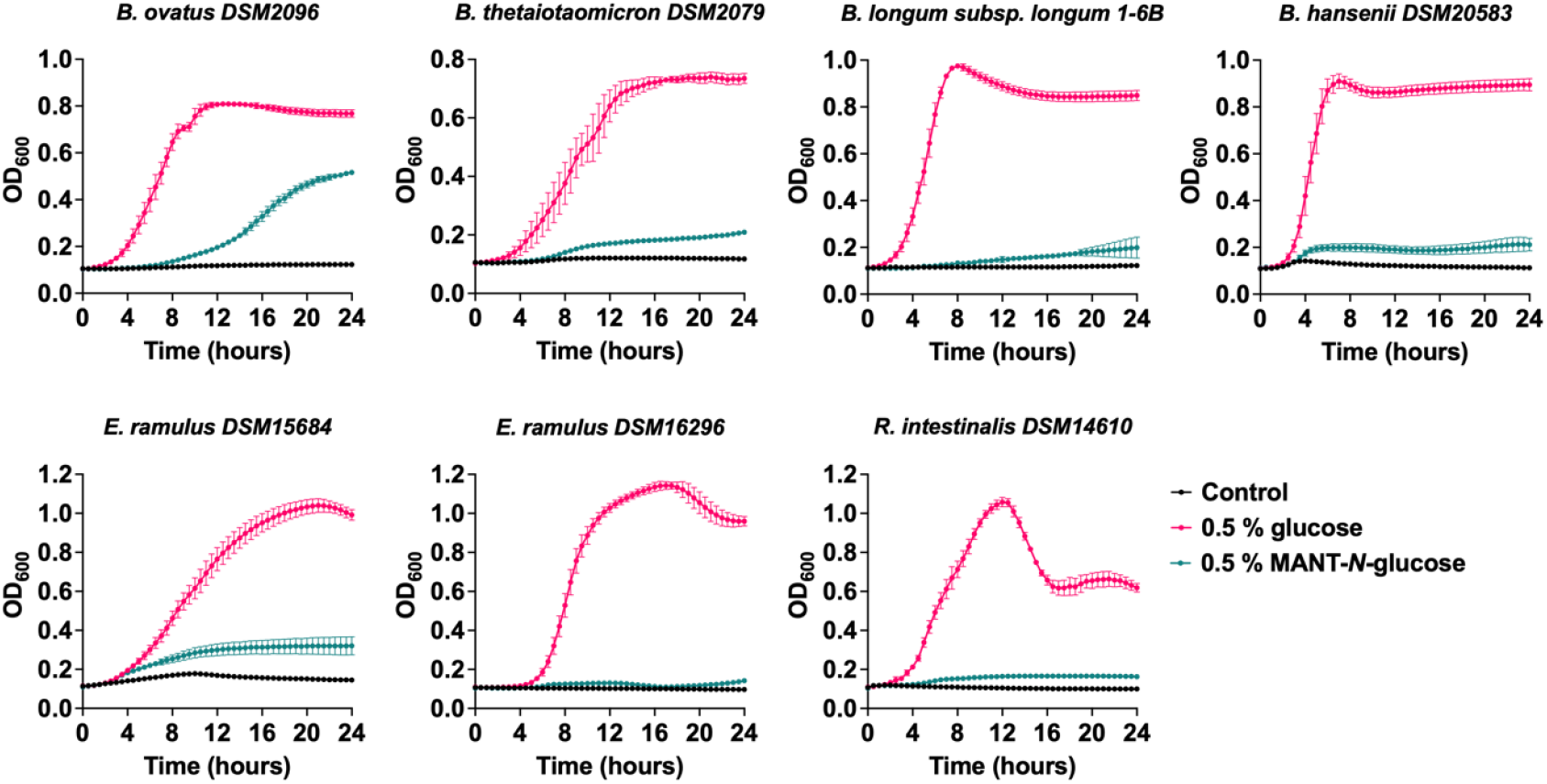
Growth curves of the seven intestinal strains supplemented with MANT-*N*-glucose, glucose, or no carbon source (control), respectively. Carbon sources (% (w/v)).

### The discovery of an *N*-glucopyranoside-hydrolysing Bgl from *Zea mays*

An in-house Bgl library (Supplementary Table 1) was screened against MANT-*N*-glucose using lysates from Bgl-expressing *E. coli*. Following an overnight reaction, it appeared that one reaction mixture had a slightly higher abundance of MANT compared to the other reaction mixtures and the control (Supplementary Figure 2). This reaction mixture contained the lysate of *E. coli*, which produced the Bgl, *Zm*Glu1 (Brzobohatý et al., 1993). The observed activity was validated by incubating the purified *Zm*Glu1 with MANT-*N*-glucose, where a significant accumulation of the aglycon, MANT, was observed compared to the control (Figure 2A, Supplementary Figure 3).

**Figure 2.**
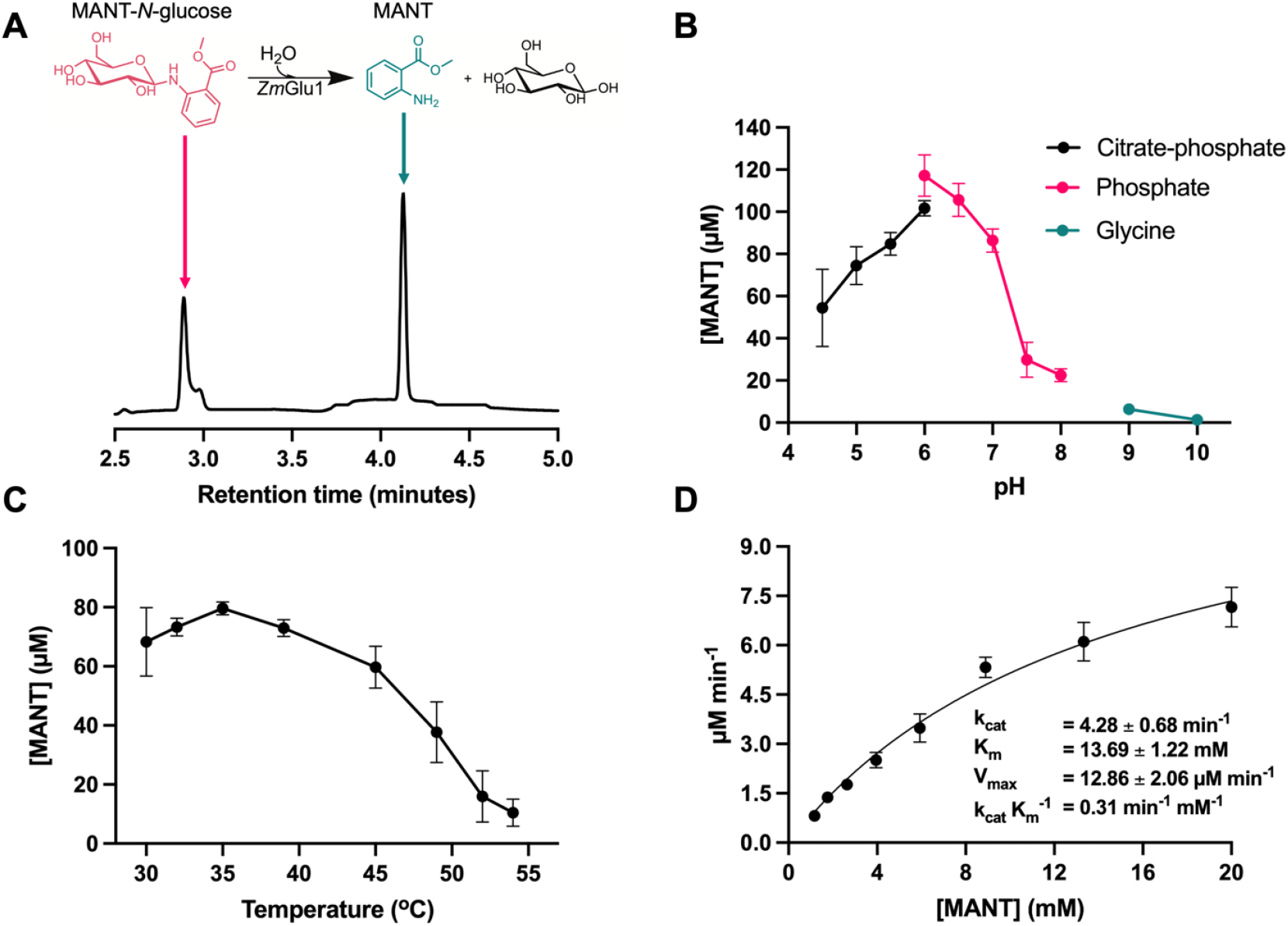
(A) The reaction equation for ZmGlu1-catalysed hydrolysis of MANT-*N*-glucose, along with a chromatogram of the overnight reaction using purified *Zm*Glu1 in citrate-phosphate buffer (pH 6.0). Peaks corresponding to MANT-*N*-glucose and MANT are indicated with arrows. (B) Determination of optimal buffer conditions. (C) Identification of the optimal reaction temperature. (D) Fitted Michaelis-Menten plot using MANT-*N*-glucose as the substrate along with the calculated kinetic properties.

**Figure 3.**
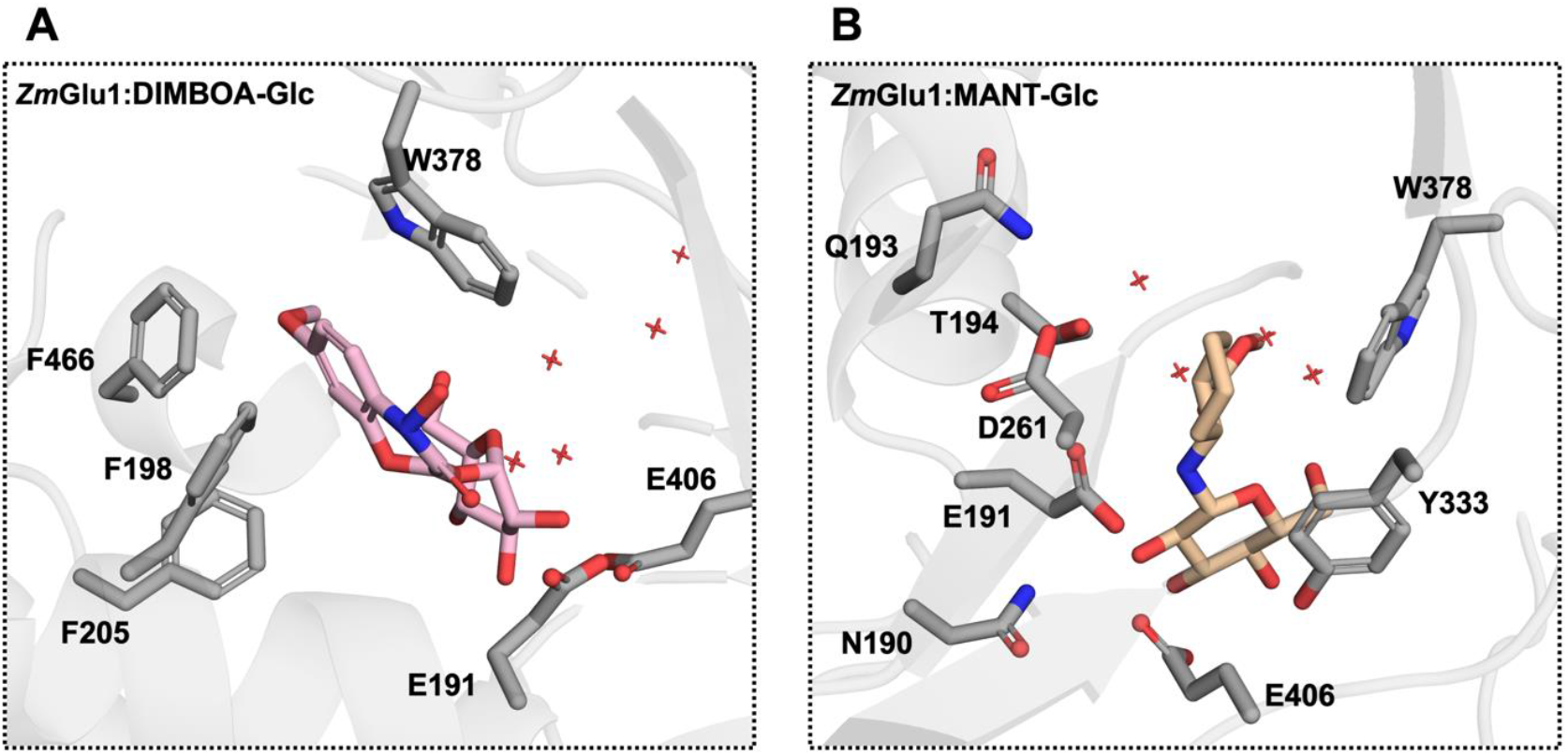
(A) DIMBOA-glucoside in *Zm*Glu1 (PDB: 1E56) with the two catalytic residues (E191 and E406) and the key residues in DIMBO-glucoside recognition. Water molecules within 10 Å of the anomeric carbon are also illustrated. These water molecules are superimposed onto the predicted *Zm*Glu1:MANT-*N*-glucose model (B). Catalytic residues and key residues in MANT-*N*-glucose recognition are illustrated along with the polar region of the active site. DIMBOA-Glc, DIMBOA-glucoside; MANT-Glc, MANT-*N*-glucose

ZmGlu1’s natural substrate is the inactive glycoside form of the antimicrobial compound, DIMBOA. It has also been shown to play a role in cytokinin release during plant growth and development (Brzobohatý et al., 1993; Gu et al., 2006). With the addition of MANT-*N*-glucose to its substrate repertoire, *Zm*Glu1’s role may extend further than previously hypothesised. Furthermore, given the small size of the Bgl library tested in this study, it is likely that plenty of other GHs with small-molecule *N*-glycopyranoside-hydrolytic activity exist in nature. However, their existence might not be described due to the significantly smaller abundance of these compounds in nature compared to the corresponding *O*-glycopyranosides (Xie et al., 2017).

### Characterisation of Bgl-catalysed hydrolysis of MANT-*N*-glucose

The *Zm*Glu1-catalysed hydrolysis of MANT-*N*-glucose was characterised. First, the optimal buffer conditions were determined, with the reaction exhibiting maximum activity at pH 6.0 in phosphate buffer (Figure 2B). This is consistent with previous reports for *Zm*Glu1 (Esen, 1992). The identical pH optima observed for ZmGlu1-catalysed hydrolysis of both *O*- and *N*-glycosides suggest that the protonation state of the two catalytic residues is critical for catalysis (Lundemo et al., 2017; McIntosh et al., 1996), supporting the notion of a shared reaction mechanism for both substrate types.

The optimal reaction temperature was determined to be 35 °C (Figure 2C), contradicting previous reports of an optimum near 50 °C (Esen, 1992). This discrepancy may be attributed to the enzyme’s reduced stability above 35 °C (Esen, 1992), which, together with the low reaction rate necessitating prolonged incubation, likely accounts for the observed difference.

The kinetic constants were determined (Figure 2D). Relative to previously reported parameters for ZmGlu1 with DIMBOA-glucoside (K_m_ = 0.098 mM, k_cat_ = 29 s^−1^) (Cicek et al., 2000) and the artificial substrate 4-nitrophenyl β-D-glucopyranoside (K_m_ = 0.38 mM, k_cat_ = 24 s^−1^) (Cicek et al., 2000; Verdoucq et al., 2003), ZmGlu1-catalysed hydrolysis of MANT-*N*-glucose was markedly slower and required higher substrate concentrations to achieve saturation. Potential explanations for this difference are addressed in the following section.

### Structural modelling of the *Zm*Glu1:MANT-*N*-glucose complex

A structural model of the ZmGlu1:MANT-*N*-glucose complex was generated using the prediction tool Chai-1 (Boitreaud et al., 2024). Across all five output structures, MANT-*N*-glucose was consistently positioned in the same orientation (Supplementary Figure 4). Comparison with the crystal structure of ZmGlu1 bound to DIMBOA-glucoside (PDB: 1E56) (Czjzek et al., 2000) revealed that the binding coordinates align closely (Supplementary Figure 4). Notably, the anomeric carbon and the *N*-glycosidic linkage are located adjacent to the two catalytic glutamates (E191 and E406), further suggesting that these residues may play a central role in *N*-glycoside hydrolysis (Figure 3).

By comparing the complex structures of *Zm*Glu1:DIMBOA-glucoside and *Zm*Glu1:MANT-*N*-glucose, we hypothesise the underlying reason for their distinct kinetic constants. First, the placement of the glucose moiety aligns in the two structures, offering similar intermolecular interactions. Hence, this portion of the structure will not be considered. In the *Zm*Glu1:DIMBOA-glucoside complex, the aglycon moiety is sandwiched between W378 and F198, F205, and F466, which can interact with DIMBOA-glucoside via pi-pi stacking and hydrophobic interactions (Figure 3A). The aglycon part of MANT-*N*-glucose is positioned slightly deeper in the pocket compared to DIMBOA-glucoside, with Y333 and W378 potentially facilitating intermolecular interactions (Figure 3B). However, we also observe that the benzene moiety of MANT-*N*-glucose is placed in a polar region of *Zm*Glu1 (Figure 3B), potentially causing unfavourable electrostatic interactions.

Furthermore, the crystal structure of the ZmGlu1:DIMBOA-glucoside complex shows the placement of water molecules within the active site (Figure 3A). When these water molecules are superimposed onto the ZmGlu1:MANT-*N*-glucose complex, positions 3–6 of the benzene ring are found to occupy the same space. This indicates that binding of MANT-*N*-glucose displaces these water molecules, possibly resulting in enthalpic and/or entropic penalties unless compensated by stabilising interactions. To test this idea, further studies are needed. Possible methods include site-directed mutagenesis of the relevant residues and *in silico* analyses, such as molecular dynamics and QM/MM simulations. These approaches could also help elucidate the reaction mechanism and assess how stabilisation of the *N*-glycoside transition state affects the reaction rate compared to *O*-glycosides.

## Conclusion

In conclusion, we have reported the discovery of the first Bgl capable of catalysing the hydrolysis of an *N*-glucopyranoside, MANT-*N*-glucose. The reaction was characterised, and its distinct kinetic properties, compared to the natural reaction, were discussed using structural models. By presenting these findings, we aim to encourage further research into Bgl-catalysed hydrolysis of *N*-glycosides, thereby advancing mechanistic understanding and supporting the identification of additional *N*-linked substrates of Bgls.

## Acknowledgements

We gratefully acknowledge financial support from the Novo Nordisk Foundation (grant NNF20CC0035580 to D.H.W.) and the Independent Research Fund Denmark (Natural Sciences, FNU, Research Project 2 grant 1026-00386B to M.A.H.). We also thank Michael Jakob Pichler for his assistance with strain selection and initial training.

